# Assessing the need for temporary fishing closures to support sustainability for a small-scale octopus fishery

**DOI:** 10.1101/2023.04.28.537864

**Authors:** Sophie Wulfing, Ahilya Sudarshan Kadba, Mez Baker-Médard, Easton R. White

## Abstract

The blue octopus (*Octopus cyanea*) fishery off the southwest coast of Madagascar is important for coastal communities. This fishery is a key economic resource for the local community as blue octopus catch is sold by local fishers to international and local export markets. Thus, it is important to monitor and evaluate the status of octopus to ensure its sustainability. One common octopus management approach is through the use of temporary spatial closures. Models can be a useful support tool to evaluate the status of a population and assess different possible management strategies. To better understand the biology and assess the sustainability of blue octopus, we parameterize a Levkovitch population matrix model using existing catch data. We found that the octopus population was experiencing a 1.8% decline per month at the time of data collection in 2006. However, since 2006, a number of management practices, including temporary closures lasting several weeks to several months have been implemented successfully. In line with these efforts, our model indicates that the fishery has likely been sustained since 2006 due to these annual closures. Our model provides support to the idea that temporary closures have restored this population and that temporary closures provide flexibility in management strategies that local communities can tailor to their economic and social needs. In addition, we were able to estimate several important life history metrics, such as time in each stage, stable stage distribution, reproductive value, and per stage survivability, that can be used in future work. Collectively, our study provides insight into the biology of blue octopus as well as demonstrate how temporary closures can be an effective conservation strategy due to the wide range of implementation options.

## 2 INTRODUCTION

Worldwide, an estimated 58.5 to 60.2 million people make their livelihood in small-scale fisheries, a subsector in which 90 to 95% of fish is distributed for local consumption, making these marine products a vital source of nutrition for these communities (FAO, 2022; FAO et al., 2023). The southwest region of Madagascar is one such area where subsistence fishing is an essential component to the diet and income of the local community. The ocean environment off the southwest coast of Madagascar is home to a wide variety of marine life, as extensive tidal flats, seagrass beds, and coral reefs are all prominent biomes in the area. In fact, Madagascar has been calculated as a country that would benefit greatly from marine conservation given its economic reliance on marine harvests and the fact that it is a refuge to many marine species (Laroche et al., 1997). In the early 2000’s, however, Madagascar’s octopus fishery began to move from local, subsistence fishing to also selling catch to export markets (Humber et al., 2006). There is evidence that up to 75% of all fish caught in select villages is now sold to outside entities for international export (Baker-Médard, 2017).

Locally-Managed Marine Areas (LMMAs) are defined as coastal and near-shore fisheries in which resources are managed almost entirely by local communities and fishery stakeholders that live in the region. Because management is conducted by those directly affected by the fishery, goals typically include maintaining the livelihood and economic and cultural goals of the local community along with environmental goals (Govan, 2010). LMMAs have grown in popularity among conservationists in small scale fisheries due to this empowerment of local fishers. Because of this, LMMAs tend to have greater local participation and compliance from stakeholders when compared to top-down regulation from governing bodies (Katikiro et al., 2015). LMMAs have been shown to improve both fisheries and fisher livelihoods in Kenya (Kawaka et al., 2017), Pacific Islands (Govan, 2010), and in Madagascar (Mayol, 2013). In Madagascar, the use of LMMAs has increased significantly since 2004, which has resulted in increased landings and Catch Per Unit Effort for local fishers (Benbow & Harris, 2011; Gilchrist et al., 2020). In order to protect fishing resources, these LMMAs instituted various conservation programs such as bans on certain types of fishing gear, implemented seasonal fishing regulations, and criminalized the harvest of endangered species.

One commonly used conservation strategy in LMMAs in Madagascar are seasonal closures. These types of reserves have a long history of use and have been seen to successfully rehabilitate stocks (Camp et al., 2015; Gnanalingam & Hepburn, 2015). For example, seasonal closures have been shown to be an effective conservation strategy in increasing biomass the Atlantic sea scallop (*Placopecten magellanicus*) fishery in the United States (Bethoney & Cleaver, 2019), restored natural trophic interactions in coral reef fisheries in Kenya (McClanahan, 2008), and successfully restored the striped marlin (*Kajikia audax*) stocks in Baja California (Jensen et al., 2010). This method is flexible, logistically simple for fishers and managers to understand, and mitigates the financial loss from the fishery that can be seen with permanent closures (Nowlis, 2000; Humber et al., 2006; Cohen & Foale, 2013; Camp et al., 2015; Gnanalingam & Hepburn, 2015; Oliver et al., 2015).

Octopus are a vital part of many ocean ecosystems and, compared to other fisheries, have a unique life history that can lead to distinct and variable population dynamics. Cephalopods act as both predators and prey in an ecosystem (Rodhouse & Nigmatullin, 1996; Santos et al., 2001; Vase et al., 2021), situating them in a key role in food webs. Further, their abundance varies drastically with a wide range of ocean conditions including sea surface and bottom temperature, salinity, currents, and sediment type (Catalán et al., 2006; Ibáñez et al., 2019; Van Nieuwenhove et al., 2019). Compared to other exploited marine organisms, octopus have a short lifespan coupled with a fast reproduction rate and high fecundity which makes their populations more responsive to fishing pressures (Langley, 2005; Humber et al., 2006). Increased fishing pressure due to globalization of the blue octopus in 2003 has since added significant fishing pressure to Madagascar’s blue octopus populations and yield from this fishery subsequently decreased in regions of this island such as the southwest region of Toliara (Langley, 2005; Humber et al., 2006). However, previous temporary closures on the fishery resulted increases in octopus, possibly indicating that this fishery has the ability to recover when fishing pressure is decreased (Humber et al., 2006; Katsanevakis & Verriopoulos, 2006; Benbow et al., 2014). However, right after reopening, catch began to decline again, which has been attributed to heavy fishing pressure right after reopening (Humber et al., 2006; Benbow et al., 2014; Oliver et al., 2015). Octopus populations are therefore sensitive to both the increase and alleviation of fishing pressure and understanding their biology and how these population dynamics will react to changes in fishing pressure is a key component to effective conservation of this resource.

*Octopus cyanea*, or blue octopus, is the most abundant cephalopod species in the western Indian Ocean and is caught in about 95% of local landings in Madagascar (Humber et al., 2006; Oliver et al., 2015). Like other cephalopod species, very little is known about their life history including natural death rate, larval survivability, and how much time this species remains in each stage of maturity. Further, age is difficult to determine from size alone as they have variable growth rates up to maturity (Wells & Wells, 1970; Heukelem, 1976; Herwig et al., 2012; Raberinary & Benbow, 2012). The *O. cyanea* that live in the southwest region of Madagascar have been shown to be genetically distinct from those outside of Madagascar (Van Nieuwenhove et al., 2019). This is because the ocean currents in the Channel are comprised primarily of eddies with very little through-flow across the Channel (Schott & McCreary, 2001; Lutjeharms et al., 2012; Hancke et al., 2014). As larval dispersion is primarily controlled by ocean currents, and *O. cyanea* does not migrate across long distances, this shows that the *O. cyanea* in Madagascar where the data was collected can be considered a distinct population (Van Nieuwenhove et al., 2019).

Size limits have been shown to be effective methods of conservation of species like *Octopus cyanea* that are harvested before maturity, and are restrictions that are easy to understand and implement in small scale fisheries (Nowlis, 2000). However, even though this is a conservation strategy often implemented in octopus fisheries, it has been shown to be less effective than instituting an overall cap on fishing effort, such as effort rotation or limiting the number if fishers (Emery et al., 2016). To protect this species, size limits have been imposed on blue octopus catch in Madagascar, but these regulations are difficult in practice, as the fishing method used to harvest octopus involves spearing the octopus’s den and extracting the octopus from the den. Blue octopus therefore typically die before size can be assessed, so octopus too small for market sale are typically harvested for household consumption (Humber et al., 2006). Further, the relationship between size and maturity stage is not strongly correlated (Raberinary & Benbow, 2012) and as a result, size restrictions wouldn’t necessarily protect the individuals ready to reproduce and would be difficult to implement in the field both due to the biology of *O. cyanea* and the characteristics of this small scale fishery. Therefore, temporary closures have been shown to be a more practical method of octopus conservation in that they can replenish stocks while maintaining fisher income (Benbow et al., 2014). Temporary closures provide many options for their duration and intensity (in other words, how much fishing can occur during a closure). However, this requires a deeper understanding of the biology and population characteristics of *O. cyanea* in this fishery in order to be properly instituted. Instituting effective temporary closures in octopus fisheries can be difficult due to their short lifespan, high mortality, and sensitivity to environmental conditions (Catalán et al., 2006; Emery et al., 2016; Ibáñez et al., 2019; Van Nieuwenhove et al., 2019). Lack of field data and difficulty of enforcement has also been a challenge in octopus fisheries, especially in Madagascar (Emery et al., 2016; Benbow et al., 2014). This indicates that a thorough understanding of the life history of *O. cyanea* and the harvest methods employed by fishers is necessary to enact meaningful fishing restrictions.

Ever since 2004, the western Madagascar region currently institutes a yearly closure of six weeks from December 15 to January 31. In addition to the regional closure, individual villages institute their own local closures once a year, typically lasting 2-3 months. These closures do not completely restrict octopus fishing, but instead institute an area where fishing is not allowed which takes up about 25% of the fishery’s spatial extent. Therefore, some octopus harvest does occur even during one of these closures (Aina, 2009; Humber et al., 2006; Benbow & Harris, 2011; Westerman & Benbow, 2014; Oliver et al., 2015; Rocliffe & Harris, 2015, 2016).

In this paper, we have three goals: 1) we will create a theoretical estimation of the species’ life history traits in different stages of its development, 2) as well as fit a Levkovitch matrix to the data collected in 2006 on *Octopus cyanea* populations in southwestern Madagascar, and 3) given this information in 2006, we will determine the frequency and length in which these temporary closures should take place to maximize population health of the fishery and maximizing catch for the local community, and show how temporary closures can be an effective conservation strategy as well as demonstrate the numerous options available when deciding the length and intensity of closures. This study is not meant to be a current stock assessment of this fishery as local communities have taken numerous steps to conserve blue octopus since the time of data collection.

## 3 METHODS

Population matrix models are a commonly used mechanistic model to predict future population dynamics by splitting the life history of the study organism up into a Leslie Matrix (Leslie, 1945) where a population is split up into groups of ages, and a transformation matrix is applied to predict what the population makeup will be in future years. As *Octopus cyanea* has an extended larval phase and there is no existing data on the age structure of this population of octopus, we use a stage-based population matrix, otherwise known as a Lefkovitch matrix (Caswell, 2001). Here, the life history of the study organism is grouped by stages (Figure 1), where each unit of the matrix represents a distinct period of the organism’s life where it is subject to different environments, pressures, or physical attributes that would alter the survival and reproductive output at that phase, but the amount of time between each stage is variable. This would simply create different inputs for the probability of remaining in the same stage, and the growth and fecundity inputs can be based on available data. Lefkovitch matrices have not yet been applied to *O. cyanea* populations and therefore could be a useful methodology to understand the dynamics of this population in the western Indian Ocean to better inform management strategies.

**Figure 1.**
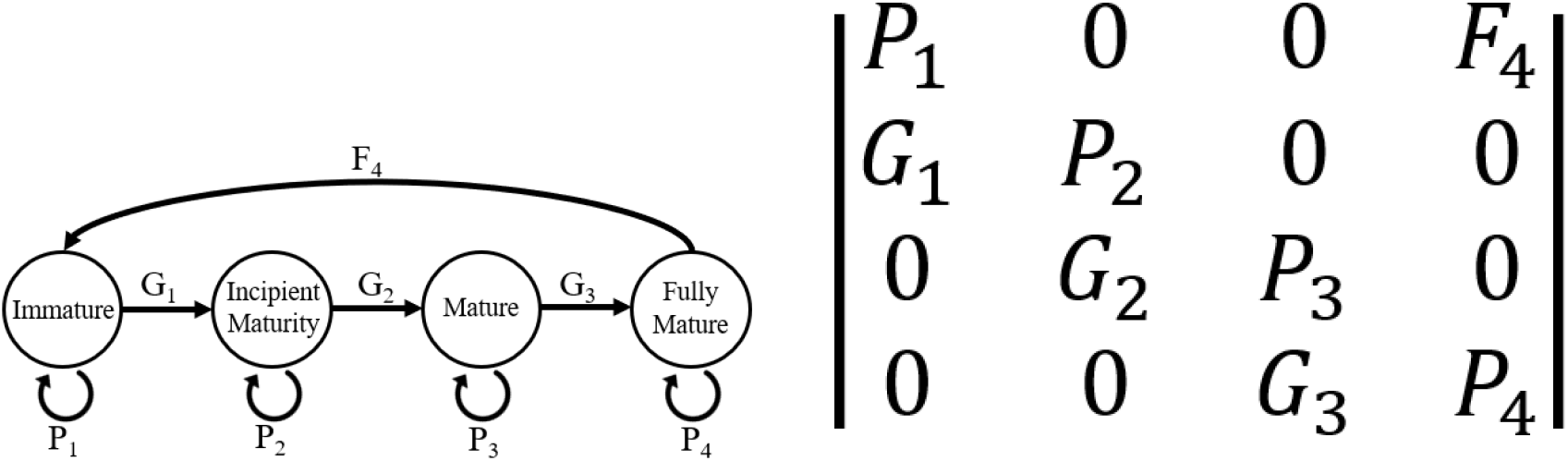
A graph representing the life history of *O. Cyanea* and the subsequent Lefkovitch Matrix where i corresponds with each of the stages of maturity (Immature, Incipient Mature, Mature, and Fully Mature individuals, respectively). *P*_*i*_ corresponds to probability of surviving and staying within a stage. *G*_*i*_ is the probability of surviving and growing to the next stage. *F*_*i*_ is the reproductive output of stage i.

### 3.1 Data

To inform our model, we use data collected by Raberinary & Benbow (2012) from landings ranging from the villages of Ampasilava in the south to Andragnombala in the north which spans about 30 kilometers of coastline. These villages are located along the Mozambique channel, where a lack of through current and prevalence of eddies results in a genetically distinct population of *O. cyanea* (Van Nieuwenhove et al., 2019). In these villages, fishers usually fish along both reef flats and deeper barrier reefs. Fishers bring catch onshore either for household consumption or to sell to buyers for international export. This study collected landing data from February 2005 to February 2006 through daily surveying fishers as they landed onshore within a two hour window. They separated each octopus into five age classes: immature, incipient maturity, maturity, full maturity, and post laying. In this paper we omit stage five, post laying, from this model as blue octopus only brood once, and stage five individuals therefore do not contribute to population growth. They recorded octopus weight, weight and length of gonads, sex, and a visual assessment of maturity class. A subsample of octopus were also collected for octopus length, and laboratory assessment of gonads for a confirmation of maturity class. They gathered this data on a total of 3,253 octopuses, and for the purposes of this study, we model from the 1,578 females collected. Despite there being no standardization for catch effort being available for this dataset, no other maturity stage study has been conducted on this population of *O. cyanea* and is therefore the best available data to fit a Lefkovitch matrix.

### 3.2 Model Parameterization

In order to parameterize this model, we use Wood’s Quadratic Programming method (Caswell, 2001). Other methods required longer time series than were available to us, were extremely sensitive to noise in the data, or simply resulted in matrices that had no reasonable biological interpretation (Caswell, 2001). One strength of Woods Quadratic Programming is it allows for constraining parameters to be within certain ranges. For example, we can constrain all parameters to be greater than zero, place zeros in the solution matrix to reflect *Octopus cyanea* biology, and ensure that all *P*_*i*_ and *G*_*i*_ parameters don’t add up to more than 1, which would imply that individuals in stage i are somehow multiplying themselves. The matrices then become quadratic equations that are solved through sum of squares minimization while also remaining within these constraints. We estimate a preliminary stage-based matrix model (Figure 2) based on Raberinary and Benbow (2012) data and calculated using the quadprog package in R (Turlach & Weingessel, 2019). We assessed model estimates by comparing life history values inferred from the matrix with existing literature on *O. cyanea* life history (Table 1). As all of our values calculated from the matrix fall within the known attributes of this species, we are confident that this model gave an accurate mechanistic description for this population’s underlying dynamics.

**Table 1:** Existing research and information on the per-stage duration of *O. cyanea*. All existing estimates are from Heukelem (1973), Heukelem (1976), Guard & Mgaya (2003), Humber et al. (2006), Aina (2009). Note: Heukelem (1976) estimate the time to maturity to be 10-13 months (i.e. stages 1-3 combined). Equations used to estimate metrics from this Lefkovitch Matrix are outlined in Barot et al. (2002).

| Stage | Existing Estimated Duration | Estimate from Lefkovitch Matrix (Months) | Standard Deviation of Estimate (Months) |
| --- | --- | --- | --- |
| Egg | 20-35 days | NA | NA |
| Larval | 28-56 days | NA | NA |
| 1: Immature | No existing estimate | 2.699666 | 2.1420858 |
| 2: Incipient Maturity | No existing estimate | 1.474724 | 0.8367118 |
| 3: Mature | No existing estimate | 1.646790 | 1.0320502 |
| 4: Fully Mature | No existing estimate | 1.494651 | 0.8598431 |
| 5: Post Laying | 45-61 days | NA | NA |
| Post Larval Phase (Stage 1-5) | 9-18 months | NA | NA |

**Figure 2.**
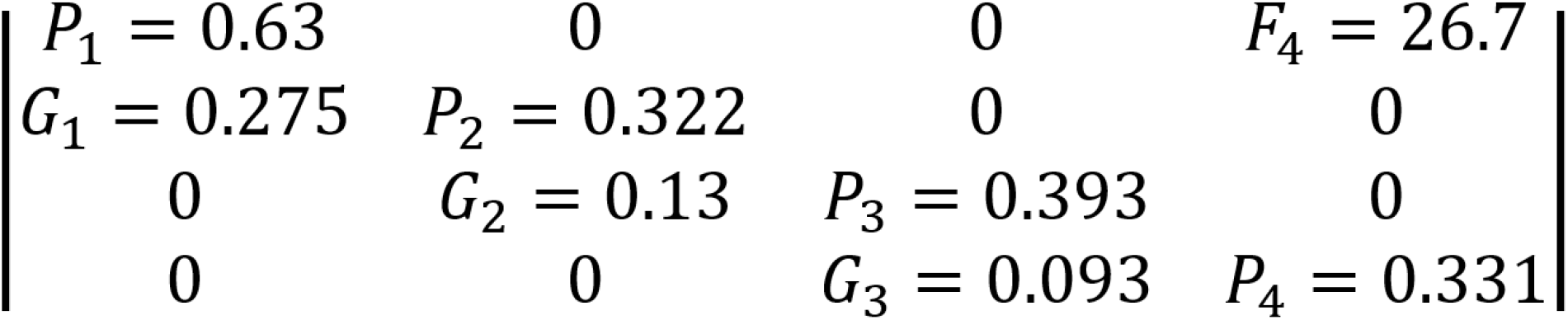
Stage-based population matrix calculated using Wood’s quadratic programming method and parameterized using data from Raberinary and Benbow (2012).

### 3.3 Model Analysis

Eigenvalues (*λ*) are calculated from the matrix and future populations can be predicted by multiplying an initial population vector to incrementally higher powers of our matrix where the power of the matrix corresponds to the time length of the projection. The initial population vector used is the blue octopus data collected in the final month of data collection from Raberinary & Benbow (2012). This month of data is not included in the parameterization of the model as it occurred after a temporary closure that was being tested at the time. We perform sensitivity analysis on the population matrix and eigenvalues using the r package popbio (Stubben & Milligan, 2007). Further, as all of the parameters are scaled to a value between 0 and 1 except *F*_4_, the different order of magnitude of these parameters have a lower proportional effect on the eigenvalue than *F*_4_. To address this, we also conduct elasticity analysis using the popbio package (Stubben & Milligan, 2007). This allows us to identify the groups within this octopus population whose protection will most benefit population growth, essentially creating focus points of conservation. The results of sensitivity and elasticity analysis are included in the supplementary material. Other life history traits that can be calculated from this matrix are stable stage distribution, reproductive value of each stage, and per-stage survivability. We also use the R package Rage (Jones et al., 2021) to calculate the age in each stage, life expectancy and longevity, the age and probability of reaching maturity, and generation time of this population. We then used the Rage package in R to analyze various life history traits of this matrix, the output of which is included in the supplementary material.

In order to incorporate uncertainty in the model, we use a Monte Carlo simulation to draw 1,000 new matrix entries from a normal distribution centered on our default estimate for that matrix entry. We test different levels of variability by increasing the standard deviation used in the random draws for each entry. The maximum standard deviation tested was 0.03 because higher standard deviations resulted in matrix entries that were too high to have realistic biological interpretations. We then calculate and evaluate the resulting dominant eigenvalues for the new matrices.

Finally, we calculate the minimum survivability increase necessary per stage in our default matrix to result in an increase of the overall population. We do this by increasing the *P*_*i*_ and *G*_*i*_ parameters by increasing percentages in each stage i until the overall eigenvalue (*λ*) became greater than one.

### 3.4 Management Scenarios

In order to determine optimal conservation strategies, we alter the survivability of *O. cyanea* by different rates from 0-10% survival increase of the species. 10% is the maximum survival increase used because increasing the overall survivability of matrix by more than 10% would result in some stages reaching a survivability of more than 1, implying that the stage would somehow be multiplying itself within a month timestep. We therefore limit survival increases to a maximum of 10% to stay within biologically meaningful parameters. Then, we simulate different closure scenarios for each survival increase by altering the length of annual closures by month using the final month of data collected by Raberinary & Benbow (2012) as the initial population vector, this is multiplied to higher powers of the original matrix during months that are simulated to be “open fishing” and then when a closure was simulated, the matrix with increased survival was multiplied to the population for that month. We simulated these different scenarios in order to analyze all combinations of conservation strategies that result in stable *O. cyanea* populations.

## 4 RESULTS

The resulting eigenvalue of our matrix is 0.982, indicating a population decline of 1.8% per month with fishing pressure included. The stable stage distribution (Table 2) shows that 65% of the makeup of this population is immature individuals, while actively breeding individuals (fully mature) only make up less than 1% of the naturally occurring population. However, the reproductive output per stage (Table 2) shows that on average, an individual in this fully mature population is expected to have 41 times the number of offspring as those in stage 1. Larval survivability of 0.0001328 is calculated by dividing our estimated number of larvae surviving back to stage 1 (*F*_4_) by 201,000 - the average estimated reproductive output of *O. cyanea* by (Guard, 2009). The life expectancy of this population is calculated by the Rage package to be 4.06 months with a standard deviation of 2.42 months. The calculated age of maturity is 6.82 months with probability of reaching maturation of 0.022. The longevity of this population (the amount of months for only 1% of the population to remain) is 12 months with a generation time of 7.38 months.

**Table 2:**
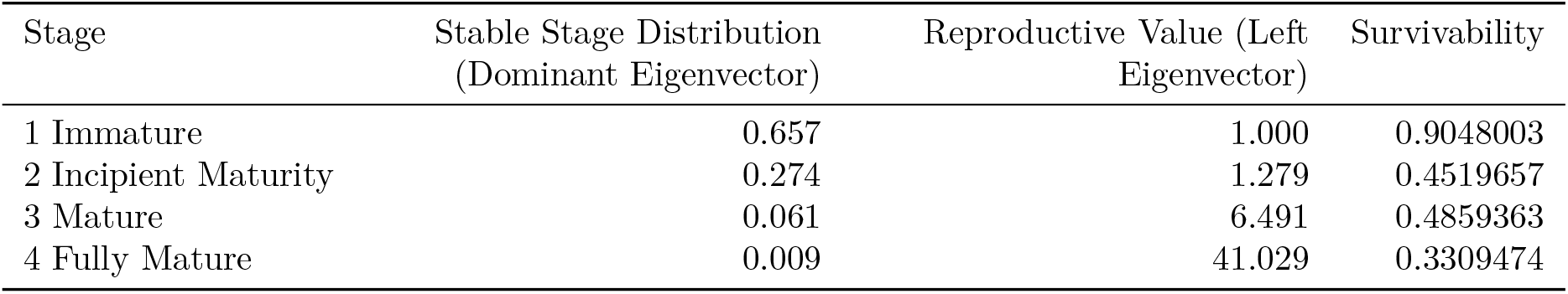
Stable stage distribution and reproductive value of each stage of this blue octopus population matrix given in Figure 2. The survivability (i.e. the proportion of individuals who survive from stage i to stage i+1) in each stage includes death rate from fishing. Stages 1-4 survivability were calculated by summing up the proportion of individuals surviving and staying within a stage every month (*P*_*i*_) and the proportion of individuals surviving and growing every month (*G*_*i*_). Larval survivability of 0.0001328 was calculated by dividing our estimated number of larvae surviving back to stage 1 (*F*_4_) by the average estimated reproductive output of *O. cyanea*.

| Stage | Stable Stage Distribution (Dominant Eigenvector) | Reproductive Value (Left Eigenvector) | Survivability |
| --- | --- | --- | --- |
| 1 Immature | 0.657 | 1.000 | 0.9048003 |
| 2 Incipient Maturity | 0.274 | 1.279 | 0.4519657 |
| 3 Mature | 0.061 | 6.491 | 0.4859363 |
| 4 Fully Mature | 0.009 | 41.029 | 0.3309474 |

The analysis of the Monte Carlo simulation shows that the resulting dominant eigenvalues are still below 1 on average, but there is a lot of variability around the mean of 0.98 (Figure 3). As the standard deviation increases in the simulation, the proportion of matrices with dominant eigenvalues above one also increases. Changing the survivability of each stage (Figure 4) shows that immature individuals (Stage 1) would need the smallest amount (5%) of survival increase in order to result in overall population growth. Stage 4, on the other hand, requires a survivability increase of 25% in order to create a viable population.

**Figure 3.**
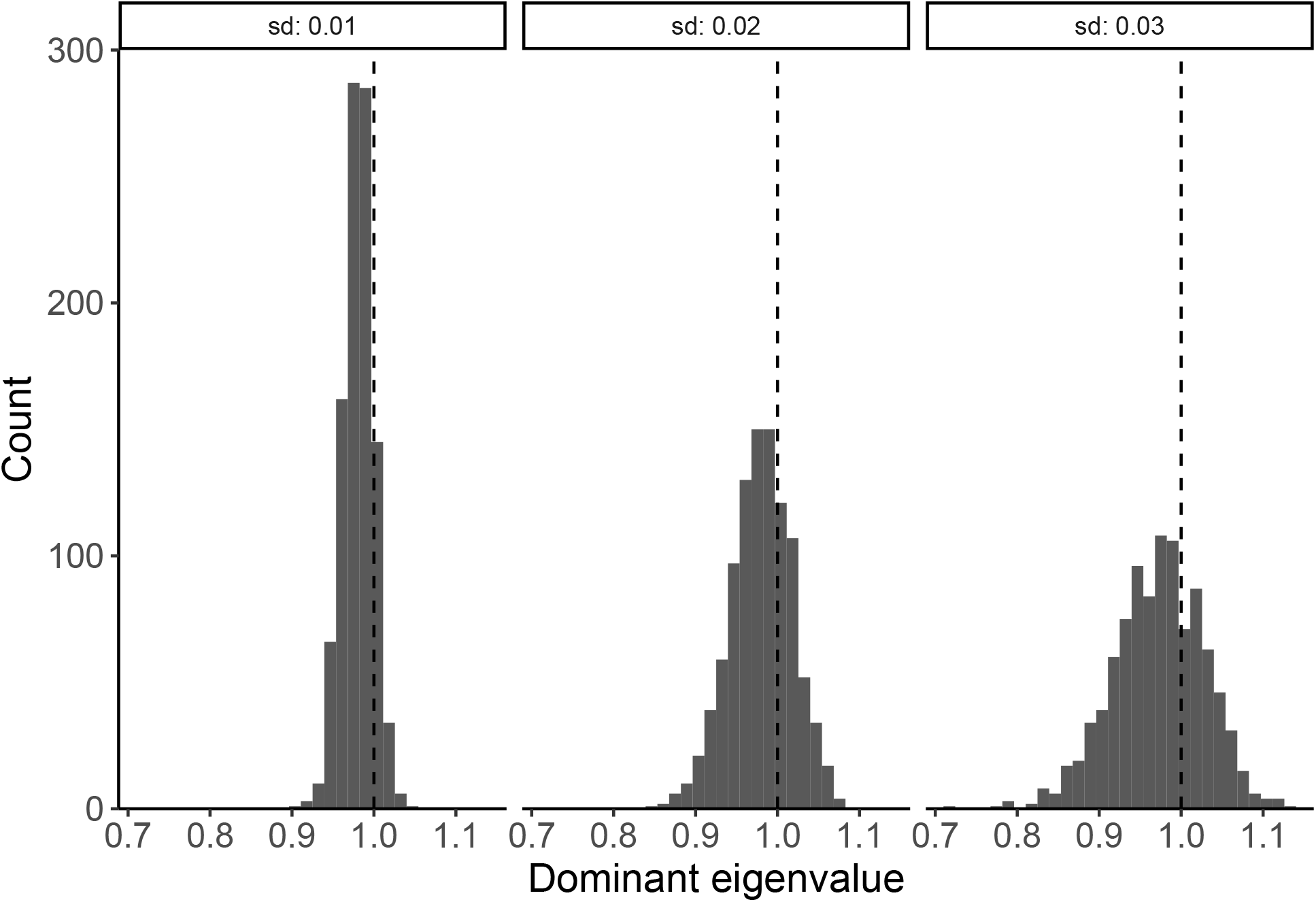
Monte Carlo simulation where matrix entries are drawn from a normal distribution centered on the default matrix at standard deviations of 0.01 (left), 0.02 (middle), and 0.03 (right). Eigenvalues are then calculated from each of the resulting matrices and plotted in a histogram after 1,000 simulations. The vertical dashed line indicates *λ* = 1, or where the eigenvalue indicates an increasing octopus population.

**Figure 4.**
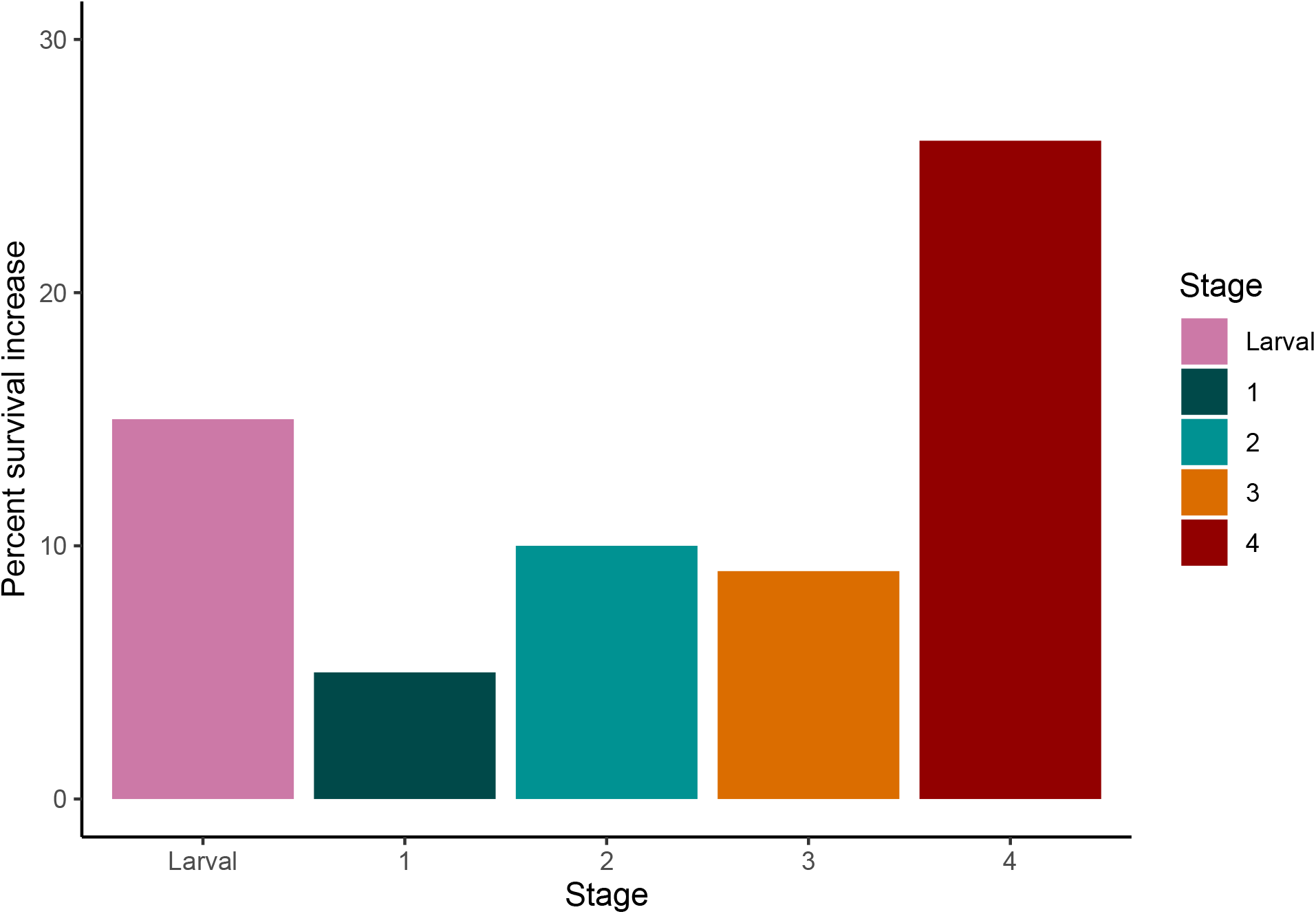
Minimum percent of per-stage survivability change needed to create population increase. Each stage was increased by higher percentages until the eigenvalue of the overall system became greater than zero.

Our analysis of different closure scenarios (Figure 5) indicates closures two months in length or shorter may be ineffective in ensuring a stable population, regardless of how much these closures decreased the death rate of the species. Further, as our baseline growth rate is close to stable (−0.0184), it took a maximum of a 7.5% increase in the survivability of the population to ensure a sustainable population when utilizing three month closures. This analysis (Figure 5) provides all the possible combinations of increased survival rates and frequency of closures that will result in a stable population. Suggested changes in overall survivability range from 2-7.5%, and the ranges of frequencies of closures span from permanent closure (every month) to once every three months.

**Figure 5.**
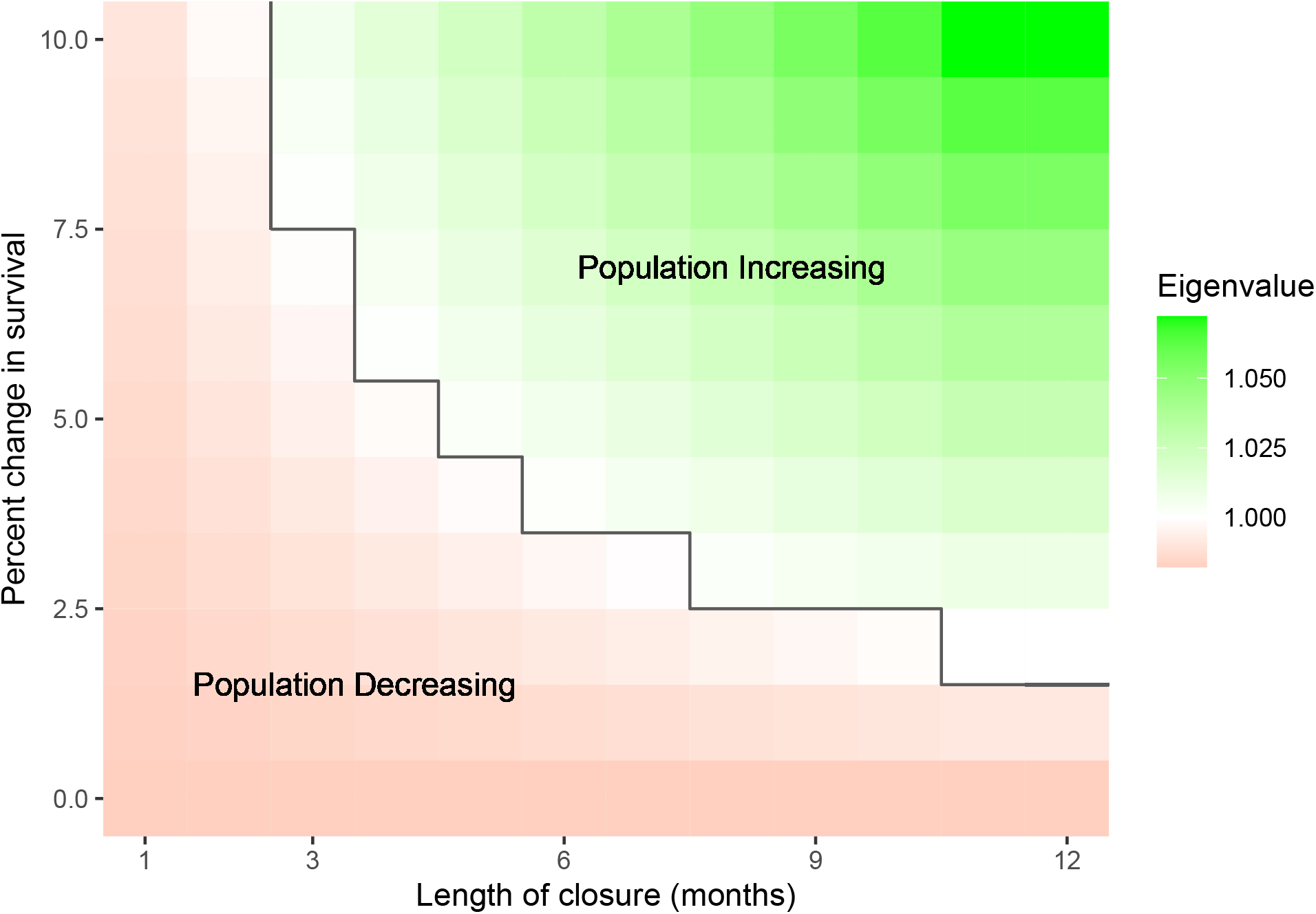
Analysis of different management scenarios. The black line separates the scenarios that succeed in sustaining the population from the scenarios that don’t. Green and white squares indicate theoretically successful management scenarios where red refers to the strategies that will not result in overall population growth.

## 5 DISCUSSION

Our model provides information about the life history of this population of *Octopus cyanea*. As each column in the matrix represents a proportion of individuals within a stage either growing or staying within a stage (with the exception of the *F*_4_ parameter), it shows a per-stage survivability estimate (Table 2) and stage duration (Table 1), life history parameters on which there has been no previous research. However, as the immature stage has a high survivability of 90.4% and a longer duration than the other stages of 2.7 months, could challenge the assumption made in the model that octopus harvest does not distinguish by size. Further, as *O. Cyanea* have an approximately one to two month larval stage (Guard & Mgaya, 2003), the fecundity parameter does not indicate the overall reproductive output of mature individuals, but the number of hatched offspring that will survive its larval stage and back the immature stage. This gives an estimation for larval survivability as female octopus have a fecundity ranging between 27,000 and 375,000 eggs (Guard, 2009), our model indicates that only an average of 26.7 individuals will survive back into immaturity, which indicates a survivability of 0.0001328. There is no other larval survivability estimation that currently exists for this species, which would be a useful further study as this could indicate a recruitment rate for this population. Further, an average lifespan of 4.06 months and an age of maturation of 6.82 months indicates that most individuals die before reaching maturation. The overall natural mortality rate of this population has been estimated to range from 0.0127 per week (0.0552 per month) to 0.0498 per week (0.2164 per month) (Roa-Ureta, 2022). However, this was not included in our model of fishery closures as the local closures do not cover the full spatial extent of the fishery, have variable spatial extents, and some fishing continues during this time, meaning some fishing mortality exists during closures (Oliver et al., 2015). Instead, we compared closures to their overall effect on the *O. cyanea* mortality rate.

Our calculated growth rate of -0.0184 and resulting population projection supports previous reports of overfishing at the time of data collection in 2006 (Humber et al., 2006; Benbow et al., 2014). With this negative growth rate, our models suggest that, without changes to management practices, the octopus population may have continued to decline. According to our model, any closure less than three months, without additional management actions, may not be effective in conserving blue octopus stocks. Uncertainty in model parameters can lead to situations where the the population would be assessed as increasing in 2006, as demonstrated by the Monte Carlo simulation (Figure 3), even though the average eigenvalue of these simulations remain below 1. This demonstrates the need for further investigation into this fishery, as octopus population dynamics can be highly variable (Humber et al., 2006), and therefore a Lefkovitch matrix may not be able to capture all of the nuances of blue octopus population dynamics. Thus, given data and model limitations, this study serves mainly to show how temporary closures are an effective tool for conservation, and caution should be taken when considering whether the octopus population was actually in decline or not in 2006. We describe this limitation more below. In general, declines in octopus populations presents an economic issue for individual fishers as their catch will become less lucrative. A successful short term closure management regimes has been shown result in economic gains from fishers in this community (Humber et al., 2006; Benbow et al., 2014; Oliver et al., 2015). Further, sale prices on opening day tend to increase as buyers are typically guaranteed larger catch (Oliver et al., 2015).

Since the time of data collection, there have been a number of important changes to fisheries management in the region (Oliver et al., 2015; Roa-Ureta, 2022). For example, temporary closures in this fishery (Oliver et al., 2015) showed that extending the regional closure beyond the conventional six weeks increased octopus catch. Further, a 2-3 month closure was suggested for this area in 2011 in order to maximize catch-per unit effort (Benbow & Harris, 2011). Benbow et al. (2014) demonstrated that a 20-week closure had similar positive effects on octopus catch when compared to a seven month closure, yet resulted in less strain on fisheries management investment than the longer seven month closure. Individual villages also institute their own closures. These closures typically span 2-3 months and restrict fishing in ∼20% of the fishery’s spatial extent, so some fishing is still allowed to occur during this time. (Rocliffe & Harris, 2015, 2016; WWF, 2017). Therefore, the changes to survivability suggested by our analysis is in relation to their overall death rate not fishing rate, indicating a need for further research on the spatial structure of this population. Our analysis of different closure scenarios suggests a range of the simplest actions needed in order to ensure sustainability of this population, and how the relationship between closure lengths and their effect on mortality rates can result in multiple different temporary closures that can successfully conserve a fishery. Thus, despite the simplicity of our model, our findings for possible closure lengths is very close to those currently practiced in Madagascar and elsewhere, and therefore suggest that temporary closure efforts in Madagascar are both necessary and have been effective in conserving this fishery. As we describe later, more realistic extensions of this model can be built to guide specific management practices.

Our analysis confirms that establishing periodic closures is an effective and commonly-used strategy in sustainable fishing practices when they are implemented deliberately (Humber et al., 2006; Oliver et al., 2015). As Madagascar has been committed to protecting its marine natural resources through increasing the number of marine parks, this study serves to highlight the effectiveness of these temporary closures and explore some of the available strategies to make population predictions and conservation strategies with limited data sources (Westlund, 2017). Implementing fishing restrictions without regard for social norms can undermine cultural practices and in turn be detrimental to both the people and fishery, and halts the dissemination of traditional ecological knowledge (Okafor-Yarwood et al., 2022). For this reason, both the Madagascar government and scientific community has found a new emphasis on studying the complex social structures within the community in question in order to more effectively conserve resources along with peoples’ livelihoods (Billé & Mermet, 2002; Baker-Médard et al., 2021). This has been shown to increase participation in conservation practices, therefore making them more effective.

The mechanistic methods used in this study allowed us to gain a baseline understanding of the growth rate and mortality of this population despite the limited data used to parameterize the model. Limitations of this study include the data collection process as this model is only parameterized using one year of data. As octopus population dynamics are extremely variable, one year of data collection may not be sufficient to get a comprehensive understanding of this population’s growth rate, as shown by the uncertainty analysis in this paper, where some simulations of uncertainty resulted in stable populations. Although this is not enough data to conduct a full stock assessment, this speaks to the utility of mechanistic modeling, where we are able to estimate population patterns and other life history traits despite this lack of data. A future study that repeats this method of data collection could rerun this same model with the updated data, and make further conclusions about the status of this fishery today. Even though data collections occurred daily within a two-hour window, catch was not standardized by effort and therefore there could be catch fluctuations between months that are not captured in the data. Further, as stage 1 had a high survival rate yet low duration, this challenges the assumption that the octopus caught are an accurate ratio of the octopus at each stage in the wild. Another shortcoming of this study is that the only available stage data for this species and region was collected in 2006, and the community of southwest Madagascar has implemented several strategies since that time to improve the sustainability of their fish stocks in the region (Humber et al., 2006; Raberinary & Benbow, 2012). Due to the time of data collection, this study does not reflect the current status of *Octopus cyanea*, nor should the findings of this study be implemented in current management decisions. In fact, more recent and comprehensive research has shown that this fishery has been fished largely sustainably from 2015 to 2017, however since then, the fishery shows evidence of operating at capacity or above (Roa-Ureta, 2022). Instead, this study outlines what biological parameters can be estimated from limited data using mechanistic modeling and show how temporary closures are not only an effective method of conservation, but also provide communities with options for effective management and these should be selected based off of the needs of stakeholders. As the community of southwest Madagascar has been involved in deciding when closures should occur and their lengths, this study serves to show the various options available (Benbow & Harris, 2011).

We made a number of simplifying assumptions in our models of the biology of the study species. For example, our models assume that all individuals within a stage are subject to the same growth and mortality rates. As this study uses data collected from a large geographic range (Raberinary & Benbow, 2012), different individuals nesting in different regions may be subject to different selective pressures. Studies on the spatial variability of this population could better inform both our model and the greater understanding of how fishing mortality of this population compares to its natural mortality. Further, this population of blue octopus has been shown to exhibit spatial variability depending on their life stage. Younger individuals tend to live in the shallow inner zone of the reef and larger individuals, who are more able to withstand stronger currents, move to deeper waters for more suitable habitats for nesting (Raberinary, 2007). Our management scenario analysis also assumes that each lifestage would be affected equally by a closure, which could be challenged by the previous result that fishers are not bringing smaller catch to landing due to the size limits. Further, explorations into uncertainty analysis of this population would help better understand the sustainability of this fishery. Despite these limitations, the data provided is the best data available for fitting a Lefkovitch matrix to this species. Future extensions of this work could include applying this method to a data rich fishery, where the conclusions of the model can be compared to empirical data. Further, valuable future research could explore the dynamics of both sexes in the population (Gerber & White, 2014) as male octopus have different growth rates and spatial dynamics (Heukelem, 1976). A better understanding of the seasonal breeding dynamics of this population of blue octopus could also give better insight into the health of this fishery (White & Hastings, 2020). Cephalopod juveniles (a key life stage in understanding future population dynamics) often have two seasonal peaks per year, indicating biannual spawning periods (Humber et al., 2006; Katsanevakis & Verriopoulos, 2006). This is related to seasonal fluctuations in temperature, as cephalopod growth is related to environmental temperature (Domain et al., 2000). However, this relationship is subject to a lot of variation (Heukelem, 1976; Herwig et al., 2012). Further, as Madagascar is a tropical climate, this trend may be different in our region of study, as suggested by Raberinary & Benbow (2012), where all life stages of O. cyanea were observed year round, suggesting continuous breeding. A better understanding the seasonality of this population could further inform when closures should take place.

## 6 CONCLUSIONS

With a short generation time, cephalopod species respond more quickly to new management strategies. As a population with highly variable population dynamics, continuous monitoring of landings, fishing effort, and where catch is found is extremely valuable in understanding the status of *Octopus cyanea* in Madagascar. This research confirms thae need for temporary closures on this fishery. Similar data has been collected by Blue Ventures on this fishery since 2015 and shows there has been an improvement to this fishery since 2006 due to local efforts, including temporary closures. Further, this collection effort does not include maturity data which would improve the analysis of this study through incorporation of multiple years of catch data (Roa-Ureta, 2022). However, this study serves to confirm the effectiveness of these closures. Finally, as the people of southwestern Madagascar are actively taking steps to conserve the health of their fisheries, we hope that studies such as these can serve to facilitate the understanding of what options are available when choosing how and when to impose fishing restrictions. We also hope that future work can build on our models to be more realistic for this system and produce specific management guidance.

## Acknowledgements

The authors would like to thank the National Science Foundation for the funding on this project [grant number 1923707]. We would also like to thank Dr. Sophie Benbow for not only collecting the data on which paper was written, but also her help in contextualizing research and answering questions about data collection.

## Data Availability

All supplemental material and code for this project are available at https://github.com/swulfing/OCyanea. All data used to parameterize this model was collected in Raberinary & Benbow (2012)

## References

Aina, T. A. N. (2009). Management of octopus fishery off south west Madagascar.

Baker-Médard, M. (2017). Gendering Marine Conservation: The Politics of Marine Protected Areas and Fisheries Access. Society & Natural Resources, 30 (6), 723–737. 10.1080/08941920.2016.1257078

Baker-Médard, M., Gantt, C., & White, E. R. (2021). Classed conservation: Socio-economic drivers of participation in marine resource management. Environmental Science & Policy, 124, 156–162. 10.1016/j.envsci.2021.06.007

Barot, S., Gignoux, J., & Legendre, S. (2002). Stage-classified matrix models and age estimates. Oikos, 96 (1), 56–61. 10.1034/j.1600-0706.2002.960106.x

Benbow, S., & Harris, A. (2011). Managing Madagascar’s octopus fisheries. Proceedingsof the workshop on Octopuscyanea fisheries, 5-6 April 2011, Toliara. Blue Ventures Conservation Report.

Benbow, S., Humber, F., Oliver, T., Oleson, K., Raberinary, D., Nadon, M., Ratsimbazafy, H., & Harris, A. (2014). Lessons learnt from experimental temporary octopus fishing closures in south-west Madagascar: Benefits of concurrent closures. African Journal of Marine Science, 36 (1), 31–37. 10.2989/1814232X.2014.893256

Bethoney, N. D., & Cleaver, C. (2019). A Comparison of Drop Camera and Diver Survey Methods to Monitor Atlantic Sea Scallops (Placopecten magellanicus) in a Small Fishery Closure. Journal of Shellfish Research, 38 (1), 43. 10.2983/035.038.0104

Billé, R., & Mermet, L. (2002). Integrated coastal management at the regional level: Lessons from Toliary, Madagascar. Ocean & Coastal Management, 45 (1), 41–58. 10.1016/S0964-5691(02)00048-0

Camp, E. V., Poorten, B. T. van, & Walters, C. J. (2015). Evaluating Short Openings as a Management Tool to Maximize Catch-Related Utility in Catch-and-Release Fisheries. North American Journal of Fisheries Management, 35 (6), 1106–1120. 10.1080/02755947.2015.1083495

Caswell, H. (2001). Matrix population models: Construction, analysis, and interpretation. Sinauer Associates.

Catalán, I. A., Jiménez, M. T., Alconchel, J. I., Prieto, L., & Muñoz, J. L. (2006). Spatial and temporal changes of coastal demersal assemblages in the Gulf of Cadiz (SW Spain) in relation to environmental conditions. Deep Sea Research Part II: Topical Studies in Oceanography, 53 (11-13), 1402–1419. 10.1016/j.dsr2.2006.04.005

Cohen, P. J., & Foale, S. J. (2013). Sustaining small-scale fisheries with periodically harvested marine reserves. Marine Policy, 37, 278–287. 10.1016/j.marpol.2012.05.010

Domain, F., Jouffre, D., & Caverivière, A. (2000). Growth of Octopus Vulgaris from tagging in Senegalese waters. Journal of the Marine Biological Association of the United Kingdom, 80 (4), 699–705. 10.1017/S0025315400002526

Emery, T. J., Hartmann, K., & Gardner, C. (2016). Management issues and options for small scale holobenthic octopus fisheries. Ocean & Coastal Management, 120, 180–188. 10.1016/j.ocecoaman.2015.12.004

FAO. (2022). The State of World Fisheries and Aquaculture 2022. Towards Blue Transformation. FAO. 10.4060/cc0461en

FAO, Duke University, & WorldFish. (2023). Illuminating Hidden Harvests: The contributions of small-scale fisheries to sustainable developmentt – Executive summary. FAO. 10.4060/cc6062en

Gerber, L. R., & White, E. R. (2014). Two-sex matrix models in assessing population viability: When do male dynamics matterx? Journal of Applied Ecology, 51 (1), 270–278. 10.1111/1365-2664.12177

Gilchrist, H., Rocliffe, S., Anderson, L. G., & Gough, C. L. A. (2020). Reef fish biomass recovery within community-managed no take zones. Ocean & Coastal Management, 192, 105210. 10.1016/j.ocecoaman.2020.105210

Gnanalingam, G., & Hepburn, C. (2015). Flexibility in temporary fisheries closure legislation is required to maximise success. Marine Policy, 61, 39–45. 10.1016/j.marpol.2015.06.033

Govan, H. (2010). Status and potential of locally-managed marine areas in the South Pacific: Munich Personal RePEc Archive, 23828.

Guard, M. (2009). Biology and fisheries status of octopus in the Western Indian Ocean and the Suitability for marine stewardship council certification (p. 22). e United Nations Environment Programme.

Guard, M., & Mgaya, Y. D. (2003). The Artisanal Fishery for Octopus cyanea Gray in Tanzania. AMBIO: A Journal of the Human Environment, 31 (7), 528–536. 10.1579/0044-7447-31.7.528

Hancke, L., Roberts, M. J., & Ternon, J. F. (2014). Surface drifter trajectories highlight flow pathways in the Mozambique Channel. Deep Sea Research Part II: Topical Studies in Oceanography, 100, 27–37. 10.1016/j.dsr2.2013.10.014

Herwig, J. N., Depczynski, M., Roberts, J. D., Semmens, J. M., Gagliano, M., & Heyward, A. J. (2012). Using Age-Based Life History Data to Investigate the Life Cycle and Vulnerability of Octopus cyanea. PLoS ONE, 7 (8), e43679. 10.1371/journal.pone.0043679

Heukelem, W. F. V. (1973). Growth and life-span of Octopus cyanea (Mollusca: Cephalopoda)*. Journal of Zoology, 169 (3), 299–315. 10.1111/j.1469-7998.1973.tb04559.x

Heukelem, W. F. V. (1976). Growth, bioenergetics, and life-span of Octopus cyanea and Octopus maya. A Dissertation Submitted to the Graduate Division of the University of Hawaii in Partial Fulfillment of the Requirements for the Degree of Doctor of Philosophy in Zoology, 232.

Humber, F., Harris, A., Raberinary, D., & Nadon, M. (2006). Seasonal Closures of No-Take Zones to promote A Sustainable Fishery for Octopus Cyanea (Gray) in South West Madagascar. Blue Ventures Conservation Report.

Ibáñez, C. M., Braid, H. E., Carrasco, S. A., López-Córdova, D. A., Torretti, G., & Camus, P. A. (2019). Zoogeographic patterns of pelagic oceanic cephalopods along the eastern Pacific Ocean. Journal of Biogeography, 46 (6), 1260–1273. 10.1111/jbi.13588

Jensen, O. P., Ortega-Garcia, S., Martell, S. J. D., Ahrens, R. N. M., Domeier, M. L., Walters, C. J., & Kitchell, J. F. (2010). Local management of a “highly migratory species”: The effects of long-line closures and recreational catch-and-release for Baja California striped marlin fisheries. Progress in Oceanography, 86 (1-2), 176–186. 10.1016/j.pocean.2010.04.020

Jones, O. R., Barks, P., Stott, I. M., James, T. D., Levin, S. C., Petry, W. K., Capdevila, P., Che-Castaldo, J., Jackson, J., Römer, G., Schuette, C., Thomas, C. C., & Salguero-Gómez, R. (2021). Rcompadre and rage - two R packages to facilitate the use of the COMPADRE and COMADRE databases and calculation of life history traits from matrix population models. bioRxiv, 2021.04.26.441330. 10.1101/2021.04.26.441330

Katikiro, R. E., Macusi, E. D., & Ashoka Deepananda, K. H. M. (2015). Challenges facing local communities in Tanzania in realising locally-managed marine areas. Marine Policy, 51, 220–229. 10.1016/j.marpol.2014.08.004

Katsanevakis, S., & Verriopoulos, G. (2006). Seasonal population dynamics of Octopus vulgaris in the eastern Mediterranean. ICES Journal of Marine Science, 63 (1), 151–160. 10.1016/j.icesjms.2005.07.004

Kawaka, J. A., Samoilys, M. A., Murunga, M., Church, J., Abunge, C., & Maina, G. W. (2017). Developing locally managed marine areas: Lessons learnt from Kenya. Ocean & Coastal Management, 135, 1–10. 10.1016/j.ocecoaman.2016.10.013

Langley, J. (2005). The 2004-2005 census of Andavadoaka, southwest Madagascar.

Laroche, J., Razanoelisoa, J., Fauroux, E., & Rabenevanana, M. W. (1997). The reef fisheries surrounding the south-west coastal cities of Madagascar. Fisheries Management and Ecology, 4 (4), 285–299. 10.1046/j.1365-2400.1997.00051.x

Leslie, P. H. (1945). On the Use of Matrices in Certain Population Mathematics. 31.

Lutjeharms, J. R. E., Biastoch, A., Van der Werf, P. M., Ridderinkhof, H., & De Ruijter, W. P. M. (2012). On the discontinuous nature of the Mozambique Current. South African Journal of Science, 108 (1/2), 5 pages. 10.4102/sajs.v108i1/2.428

Mayol, T. (2013). Madagascar’s nascent locally managed marine area network. Madagascar Conservation & Development, 8 (2), 91–95. 10.4314/mcd.v8i2.8

McClanahan, T. R. (2008). Response of the coral reef benthos and herbivory to fishery closure management and the 1998 ENSO disturbance. Oecologia, 155 (1), 169–177. 10.1007/s00442-007-0890-0

Nowlis, J. S. (2000). Short- and long-term effects of three fishery-management tools on depleted fisheries. BULLETIN OF MARINE SCIENCE, 66 (3), 12.

Okafor-Yarwood, I., Kadagi, N. I., Belhabib, D., & Allison, E. H. (2022). Survival of the Richest, not the Fittest: How attempts to improve governance impact African small-scale marine fisheries. Marine Policy, 135, 104847. 10.1016/j.marpol.2021.104847

Oliver, T. A., Oleson, K. L. L., Ratsimbazafy, H., Raberinary, D., Benbow, S., & Harris, A. (2015). Positive Catch & Economic Benefits of Periodic Octopus Fishery Closures: Do Effective, Narrowly Targeted Actions “Catalyze” Broader Management? PLOS ONE, 10 (6), e0129075. 10.1371/journal.pone.0129075

Raberinary, D. (2007). Periode de ponte du poulpe (Octopus cyanea) D’Andavadoaka dans la region sud oest de Madagascar. Blue Ventures Conservation.

Raberinary, D., & Benbow, S. (2012). The reproductive cycle of Octopus cyanea in southwest Madagascar and implications for fisheries management. Fisheries Research, 125–126, 190–197. 10.1016/j.fishres.2012.02.025

Roa-Ureta, R. H. (2022). Stock Assessment of Octopus cyanea in the fishery of Southwest Madagascar, 2015 to 2020 (p. 26). Blue Ventures.

Rocliffe, S., & Harris, A. (2015). Scaling success in octopus fisheries management in the Western Indian Ocean. Proceedings of the workshop, 3-5 December 2014, Stone Town, Zanzibar. Blue Ventures.

Rocliffe, S., & Harris, A. (2016). Blue Ventures Report: Experiences of periodic closures in small-scale invertebrate fisheries April 2016. Blue Ventures.

Rodhouse, P. G., & Nigmatullin, M. (1996). Role as consumers. Royal Society Publishing, 20. 10.1098/rstb.1996.0090

Santos, M. B., Clarke, M. R., & Pierce, G. J. (2001). Assessing the importance of cephalopods in the diets of marine mammals and other top predators: Problems and solutions. Fisheries Research, 52 (1-2), 121–139. 10.1016/S0165-7836(01)00236-3

Schott, F. A., & McCreary, J. P. (2001). The monsoon circulation of the Indian Ocean. Progress in Oceanography, 51 (1), 1–123. 10.1016/S0079-6611(01)00083-0

Stubben, C. J., & Milligan, B. G. (2007). Estimating and analyzing demographic models using the popbio package in r. Journal of Statistical Software, 22 (11).

Turlach, B. A., & Weingessel, A. (2019). Quadprog: Functions to solve quadratic programming problems. https://CRAN.R-project.org/package=quadprog

Van Nieuwenhove, A. H. M., Ratsimbazafy, H. A., & Kochzius, M. (2019). Cryptic diversity and limited connectivity in octopuses: Recommendations for fisheries management. PLOS ONE, 14 (5), e0214748. 10.1371/journal.pone.0214748

Vase, V. K., Koya, M. K., Dash, G., Dash, S., Sreenath, K. R., Divu, D., Kumar, R., Rahangdale, S., Pradhan, R. K., Azeez, A., Sukhdhane, K. S., & Jayshree, K. G. (2021). Acetes as a Keystone Species in the Fishery and Trophic Ecosystem Along Northeastern Arabian Sea. Thalassas: An International Journal of Marine Sciences, 37 (1), 367–377. 10.1007/s41208-020-00276-y

Wells, M. J., & Wells, J. (1970). Observations on the feeding, growth rate and habits of newly settled Octopus Cyanea. Journal of Zoology, 161 (1), 65–74. 10.1111/j.1469-7998.1970.tb02170.x

Westerman, K., & Benbow, S. (2014). The Role of Women in Community-based Small-Scale Fisheries Management: The Case of the South West Madagascar Octopus Fishery. Western Indian Ocean Journal of Marine Science, 12 (2), 119–132.

Westlund, L. (Ed.). (2017). Marine protected areas: Interactions with fishery livelihoods and food security. Food and Agriculture Organization of the United Nations.

White, E. R., & Hastings, A. (2020). Seasonality in ecology: Progress and prospects in theory. Ecological Complexity, 44, 100867. 10.1016/j.ecocom.2020.100867

WWF. (2017). Abundant octopus on the Mahafaly coast. World Wildlife Fund.

