## Supplemental Material for "Assessing the need for temporary fishing closures to support sustainability for a small-scale octopus fishery"

Supplementary Material for:

### Using mechanistic models to assess temporary closure strategies for small scale fisheries

Sophie Wulfing[1*], Ahilya Sudarshan Kadba[1], Mez Baker-Médard[2], and Easton R. White[1]
[1]Department of Biological Sciences, University of New Hampshire, 03824, NH, USA

[2]Department of Environmental Studies, Middlebury College, Middlebury, VT 05753

Table 1: Data collected in Raberinary and Benbow 2012.

| T | Stage1 | Stage2 | Stage3 | Stage4 | Total |
|---|---|---|---|---|---|
| 1 | 10.40119 | 5.200594 | 0.000000 | 1.1144131 | 16.71620 |
| 2 | 76.52303 | 27.860327 | 2.228826 | 1.8573551 | 108.46954 |
| 3 | 57.57801 | 37.890045 | 1.857355 | 0.0000000 | 97.32541 |
| 4 | 40.49034 | 50.891531 | 3.343239 | 0.0000000 | 94.72511 |
| 5 | 71.69391 | 16.716196 | 8.172363 | 1.1144131 | 97.69688 |
| 6 | 121.09955 | 28.974740 | 5.572065 | 2.2288262 | 157.87519 |
| 7 | 119.98514 | 52.005944 | 6.686478 | 0.7429421 | 179.42051 |
| 8 | 78.75186 | 41.604755 | 14.487370 | 1.1144131 | 135.95840 |
| 9 | 118.87073 | 53.491828 | 14.487370 | 1.1144131 | 187.96434 |
| 10 | 119.98514 | 39.004458 | 10.772660 | 1.1144131 | 170.87667 |
| 11 | 73.55126 | 26.374443 | 4.457652 | 2.2288262 | 106.61218 |
| 12 | NA | NA | NA | NA | NA |
| 13 | 83.58098 | 111.069837 | 21.545320 | 1.1144131 | 217.31055 |

### 1 Data

Table 1 shows data used to parameterize matrix model from Raberinary and Benbow (2012). Data was extracted from Figure 7 of this paper using WebPlotDigitizer (https://automeris.io/WebPlotDigitizer/)

### 2 Stability and Elasticity analysis

Sensitivity analysis (Figure 1) showed that within each stage, the growth parameters ($G_1$ - $G_3$) had the largest effect on this growthrate compared to the parameters indicating staying within a stage ($P_1$-$P_4$). However, as all the parameters represent proportions of individuals in a stage and must necessarily be between 0 and 1 with the exception of the $F_4$ parameter, elasticity analysis provides an interpretation that weights all stages equally. The result of this analysis shows that percent changes in the fecundity metric can be as beneficial to the overall population growth as changes in the G parameters (Figure 2). Further, this analysis indicates that of all the stages, stage 1 has the most overall influence on the overall population growth.

Elasticity analysis shows that conservation of both the growth and reproductive parameters would have equal effect on the overall population growth, with the most influential parameter being the survival of stage 1 individuals. The sensitivity and elasticity analysis indicate which stages will have the greatest effect on the population if they are targeted for preservation practices. Previous research has indicated that catch size limits are very effective in preserving stocks of species with rapid growth and high death rates, but this is only if individuals do not enter the fishery until they have reached maturity (Nowlis 2000). However, the fishing method most commonly employed by the local people is spearfishing, where harvesters will search out octopus dens and spear the den to probe out the octopus (Benbow et al. 2014). Because of this, fishing method does not discriminate based on stage, this is not an applicable suggestion for conservation practices.

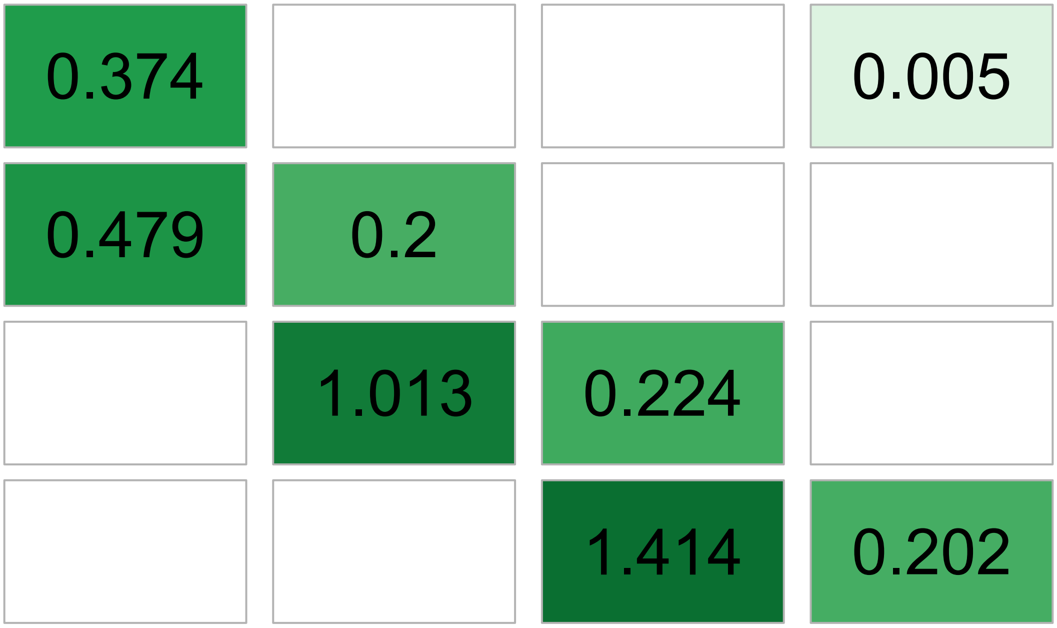


Figure 1: Sensitivity analysis of our matrix model - the change in the eigenvalue ($\lambda$) as a result of a unit change of each parameter in the model.

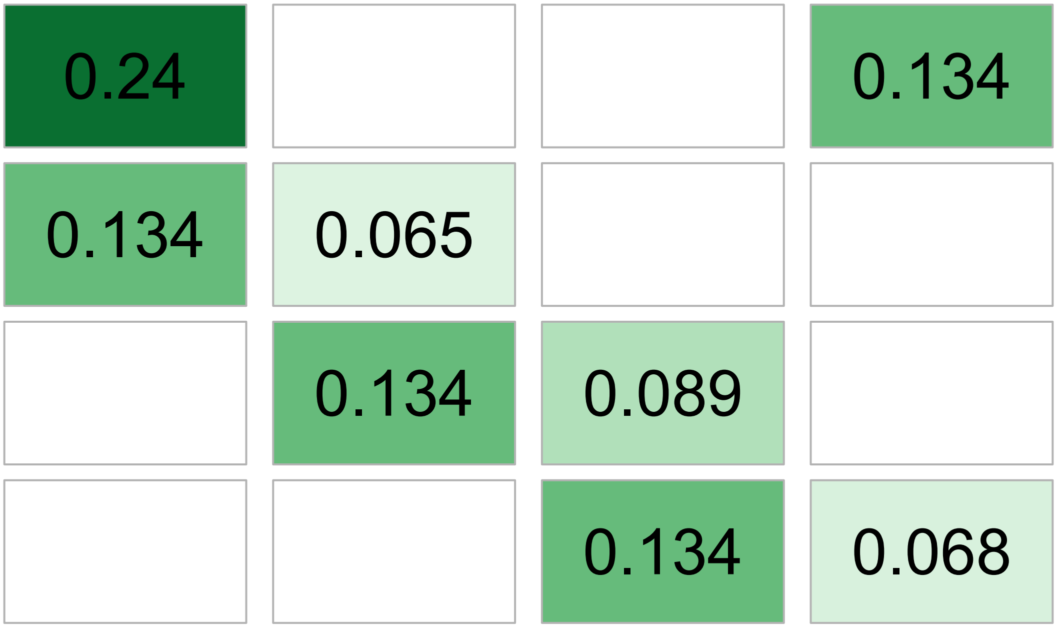


Figure 2: Elasticity analysis of our matrix model - the change in the eigenvalue ($\lambda$) as a result of a proportional change of each paramter in the model.

For this reason, even though our analysis of different stage survivabilities indicates that conserving immature individuals would be an effective tool for fostering population growth, this is not a realistic management practice in this fishery for most harvesters.

### 3 Per-stage management scenarios

#### 3.1 METHODS

Examined how increasing the chance of survival of individuals in each stage class would contribute to population health. This was achieved by isolating the growth ($G_i$) and in-stage survival ($P_i$) for each stage i. We then increased these parameters by 1% and recalculated the overall eigenvalue of the matrix. We then incorporated into different scenarios with different frequencies of fishing restrictions to examine how temporary closures on blue octopus in certain stages would affect the population.

#### 3.2 RESULTS

Our within stage analysis showed that Stage 1 needed the smallest percent increase in survival to result in overall population growth (Figure 3). Stage 4 and larval survivability would be the highest needed increase, with stage 4 needing a 25% increase and larval needing a 15% increase in order for the overall population to be stable. Further, when examining the different frequencies of fishing closures, we found that, for any scenario, no closure would be effective if it was less frequent than every other month. As exemplified by the previous analysis, closures focusing on either stage 4 or larval individuals required the most increase in survivorship and highest frequencies of closures in order to result in population growth.

#### 3.3 DISCUSSION

The results of our per-stage analysis showed that focusing on protecting individuals in stage 1 would be the most effective form of management if size could be determined before capture in this fishery. It is a common trend that individuals that survive long larval stages that have high death rates are the most valuable in terms of contributions to overall population growth. However, this is not a realistic management suggestion, as it is difficult to assess the size of octopus before catch, which is often fatal to the individual. This could suggest, however, that the establishment of aquaculture of *Octopus cyanea* could have benefits to the overall population if octopus are reared until passing this first stage of development. However, further research is needed on cephalopod aquaculture in order to be effective and reduce pollutants in the surrounding waters (Jacquet et al. 2023).

### 4 Rage package analysis

#### 4.1 Age specific Calculations

Table 2 shows age specific life-history traits of *Octopus cyanea* as calculated by the Rage package from our matrix. Expected number of offspring is reported per original cohort number.

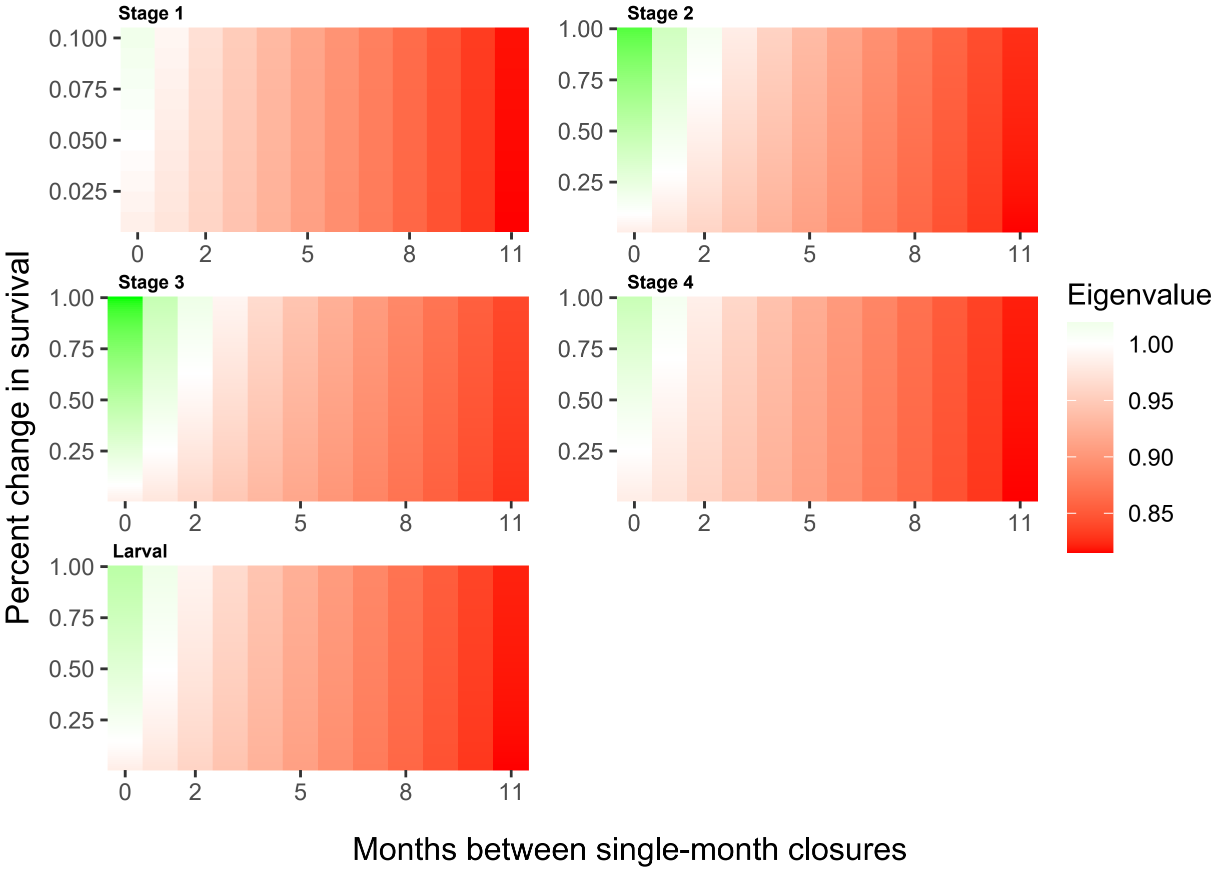


Figure 3: Different fishing scenarios based on increasing survivability of one stage.

Table 2: Life table of *O. cyanea* as calculated from our Lefkovitch matrix.

| Age (months) | Survivorship | Proportion of orginal cohort dying | Mortality hazard | Probability of death | Probability of survival | Remaining life expectancy | Per-capita reproduction rate | Expected number of offspring |
|---|---|---|---|---|---|---|---|---|
| 0 | 1.0000000 | 0.0951997 | 0.0999577 | 0.0951997 | 0.9048003 | 3.546039 | 0.0000000 | 0.0000000 |
| 1 | 0.9048003 | 0.2107642 | 0.2636471 | 0.2329401 | 0.7670599 | 2.866532 | 0.0000000 | 0.0000000 |
| 2 | 0.6940360 | 0.1996469 | 0.3359857 | 0.2876607 | 0.7123393 | 2.585199 | 0.0000000 | 0.0000000 |
| 3 | 0.4943891 | 0.1567056 | 0.3766632 | 0.3169681 | 0.6830319 | 2.427255 | 0.1801271 | 0.0890529 |
| 4 | 0.3376835 | 0.1130954 | 0.4022801 | 0.3349152 | 0.6650848 | 2.321617 | 0.4417776 | 0.1491810 |
| 5 | 0.2245882 | 0.0778436 | 0.4192659 | 0.3466059 | 0.6533941 | 2.238925 | 0.7077949 | 0.1589624 |
| 6 | 0.1467446 | 0.0520106 | 0.4307673 | 0.3544291 | 0.6455709 | 2.161373 | 0.9405289 | 0.1380175 |
| 7 | 0.0947340 | 0.0340773 | 0.4386006 | 0.3597150 | 0.6402850 | 2.073494 | 1.1279077 | 0.1068513 |
| 8 | 0.0606568 | 0.0220360 | 0.4439266 | 0.3632896 | 0.6367104 | 1.957489 | 1.2709528 | 0.0770919 |
| 9 | 0.0386208 | 0.0141236 | 0.4475285 | 0.3656983 | 0.6343017 | 1.789093 | 1.3761398 | 0.0531476 |
| 10 | 0.0244972 | 0.0089981 | 0.4499474 | 0.3673119 | 0.6326881 | 1.532303 | 1.4513443 | 0.0355539 |
| 11 | 0.0154991 | 0.0057097 | 0.4515599 | 0.3683858 | 0.6316142 | 1.131614 | 1.5039407 | 0.0233097 |
| 12 | 0.0097895 | 0.0097895 | 2.0000000 | 1.0000000 | 0.0000000 | 0.500000 | 1.5400772 | 0.0150765 |

#### References

Benbow, Sophie, F Humber, Ta Oliver, Kll Oleson, Daniel Raberinary, M Nadon, H Ratsimbazafy, and A Harris. 2014. "Lessons Learnt from Experimental Temporary Octopus Fishing Closures in South-West Madagascar: Benefits of Concurrent Closures." *African Journal of Marine Science* 36 (1): 31–37. https://doi.org/10.2989/1814232X.2014.893256.

Jacquet, Jennifer, Becca Franks, Peter Godfrey-Smith, and Walter Sánchez-Suárez. 2023. "The Case Against Octopus Farming."

Nowlis, Joshua Sladek. 2000. "Short- and Long-Term Effects of Three Fishery-Management Tools on Depleted Fisheries." *Bulletin of Marine Science* 66 (3): 12.

Raberinary, Daniel, and Sophie Benbow. 2012. "The Reproductive Cycle of Octopus Cyanea in Southwest Madagascar and Implications for Fisheries Management." *Fisheries Research* 125-126 (August): 190–97. https://doi.org/10.1016/j.fishres.2012.02.025.
